# A Multi-Omics Atlas of Sex-Specific Differences in Obstructive Hypertrophic Cardiomyopathy

**DOI:** 10.1101/2024.02.22.581621

**Authors:** Ramin Garmany, Surendra Dasari, J. Martijn Bos, Evelyn T. Kim, Katherine A. Martinez, David J. Tester, Cristobal dos Remedios, Joseph J. Maleszewski, Joseph A. Dearani, Steve R. Ommen, Jeffrey B. Geske, John R. Giudicessi, Michael J. Ackerman

## Abstract

**Background:** Hypertrophic cardiomyopathy (HCM) is a common genetic heart disease. Women with HCM tend to have a later onset but more severe disease course. However, the underlying pathobiological mechanisms for these differences remain unknown.

**Methods:** Myectomy samples from 97 patients (53 males/44 females) with symptomatic obstructive HCM and 23 control cardiac tissues were included in this study. RNA-sequencing was performed on all samples. Mass spectrometry-based proteomics and phosphoproteomics was performed on a representative subset of samples.

**Results:** The transcriptome, proteome, and phosphoproteome was similar between sexes and did not separate on PCA plotting. Overall, there were 482 differentially expressed genes (DEGs) between control females and control males while there were only 53 DEGs between HCM females and HCM males. There were 1963 DEGs between HCM females and control females compared to 1064 DEGs between HCM males and control males. Additionally, there was increased transcriptional downregulation of hypertrophy pathways in HCM females and in HCM males. HCM females had 119 differentially expressed proteins compared to control females while HCM males only had 27 compared to control males. Finally, the phosphoproteome showed females had 341 differentially phosphorylated proteins (DPPs) compared to controls while males only had 184. Interestingly, there was hypophosphorylation and inactivation of hypertrophy pathways in females but hyperphosphorylation and activation in males.

**Conclusion:** There are subtle, but biologically relevant differences in the multi-omics profile of HCM. This study provides the most comprehensive atlas of sex-specific differences in the transcriptome, proteome, and phosphoproteome present at the time of surgical myectomy for obstructive HCM.

## Introduction

Sex is a well-established biological variable impacting cardiovascular diseases^1^. Despite this awareness, health disparities between males and females are concerning with cardiovascular diseases among the leading cause of death in women^1, 2^. As such, there have been attempts to increase inclusion of women in cardiovascular clinical trials to help identify sex-specific clinical differences^2^. Nevertheless, the inclusion of women remains limited in preclinical studies with women underrepresented in basic and translational cardiovascular research^3^. Additionally, the lack of reporting of sex-specific data by separating results for males and female prevents investigation of the sex-specific differences in disease^4^.

Clinical disparities between males and females have also been in observed in the diagnosis, clinical course, and outcomes in hypertrophic cardiomyopathy (HCM)^5–9^. HCM is a genetic heart disease characterized by left ventricular hypertrophy and a diverse range of clinical presentations from an asymptomatic course to sudden cardiac arrest or heart failure^10^. Several multi-center studies have shown that females with HCM present at a later age with an increased symptom burden and higher risk of ventricular arrhythmias, heart failure, and atrial fibrillation^11–14^.

Recently, we performed one of the largest multi-omics studies on myectomy samples from patients with obstructive HCM identifying a common transcriptional and proteomic dysregulation, irrespective of genotype^15, 16^. Additionally, we found subtle differences between genotype-positive - and genotype-negative patients which might explain the clinical differences observed in these subgroups^16^. We hypothesized that, like genotype, there are molecular differences between males and females with HCM which may drive some of the observed, sex-specific clinical differences. Herein we provide, to our knowledge the first, and most comprehensive, sex-specific multi-omics profiling of myectomy tissue from patients with HCM to identify sex-specific differences in obstructive HCM.

## Methods

The Mayo Clinic IRB approved study consisted of 97 myectomy samples from patients (53 males/44 females) undergoing septal reduction therapy for symptomatic, obstructive HCM. Control samples were derived from cardiac transplant donors with a normal cardiac examination for whom suitable recipients were not found^15^. RNA-sequencing and analyses were performed on all samples as previously described^15^. Differential expression analysis was performed comparing female controls to male controls, HCM females to HCM males, HCM female to female controls, and HCM males to male controls. Transcripts with an adjusted differential expression p-value of ≤ 0.05 and an absolute log2 fold change of ≥ 1 were considered differentially expressed genes (DEGs). DEGs were subsequently inputted into Ingenuity Pathway Analysis (IPA, Qiagen, Hilden, Germany) to identify altered canonical pathways.

Pathways with a -log (B-H p-value) ≥ 1.3 were considered altered. If z-score ≥ 2 pathways were classified as upregulated, if 1 ≤ z-score ≤ 2 pathways were considered moderately upregulated, if z-score ≤ -2 pathways were considered downregulated, if -2 ≤ z-score ≤ -1 pathways were considered moderately downregulated, and otherwise directionality was undetermined.

The proteomics analysis consisted of a representative, statistically powered subset of the samples including 12 HCM males, 12 HCM females, 3 control females, and 3 control males. As previously described, samples were TMT-labeled and underwent nano-scale liquid chromatography tandem mass spectometry (nLC-MS/MS)^15^. Additionally, a fraction of each sample underwent enrichment for phosphopeptides^15^. Protein identification and quantification was performed as previously described^15^. Differential analysis was performed to compare control females to control males, HCM females to HCM males, HCM females to female controls, and HCM males to male controls. Protein groups that had an adjusted differential expression p-value ≤ 0.05 and a log2 fold change of ≥ |0.5| (where 0.0 is considered ‘no change’) were considered differentially expressed proteins (DEPs) or differentially phosphorylated proteins (DPPs). DEPs and DPPs were inputted into Ingenuity Pathway Analysis to identify altered canonical pathways. Pathways with a -log (B-H p-value) ≥ 1.3 were considered altered. If z-score ≥ 2 pathways were classified as upregulated, if 1 ≤ z-score ≤ 2 pathways were considered moderately upregulated, if z-score ≤ -2 pathways were considered downregulated, if -2 ≤ z-score ≤ -1 pathways were considered moderately downregulated, and otherwise directionality was undetermined.

## Results

### Similar Baseline Characteristics Between Males and Females

**Supplemental Table 1** includes demographic and clinical information on the females and males with HCM. Overall, all baseline, clinical characteristics as assessed at time of surgical myectomy were similar between females and males. Importantly, age at myectomy was similar between both groups thus not a confounding variable.

### Transcriptome of HCM is Similar Between Sexes

In total 19,554 transcripts were identified across the cardiac samples. As expected, from PCA analyses, the transcriptome of HCM is distinct from controls; however, there is little to no separation between males and females (**Supplemental Figure 1A**). Interestingly, we observed more variability between control males and females than HCM males and females suggesting HCM abolishes some sex-specific differences (**Supplemental Figure 1A**). Next, four different differential expression analyses were performed: control females vs. control males (**Supplemental Figure 1B**), HCM females vs. HCM males (**Supplemental Figure 1C**), HCM females vs. HCM males (**Supplemental Figure 1D**), HCM males vs. control males (**Supplemental Figure 1E**).

### HCM Abolishes Sex Specific Transcriptional Differences

In total, there were 482 DEGs between control females and control males compared to only 53 DEGs between HCM females and HCM males (**Figure 1A**). Of the 482 transcripts that differentiated sexes in the healthy tissue, only 44 (9%) were differentially expressed between HCM females and HCM males. Interestingly, there were 9 DEGs uniquely different between HCM females and HCM males, including upregulation of *Xist*_exon4, *VGLL2*, and *ESM1,* and downregulation of *PPM1E, TUBA3E*, *CORIN*, *GLP1R*, *DHRS7C*, and *CPNE5*. Pathway analysis of the 53 DEGs between HCM females and HCM males showed a total of 5 altered pathways, 4 of which were also altered in controls and only 1 (20%) pathway, Retinoate Biosynthesis I (- log[B-H p-value]=1.33), was uniquely altered in HCM (**Figure 1B**).

**Figure 1.**
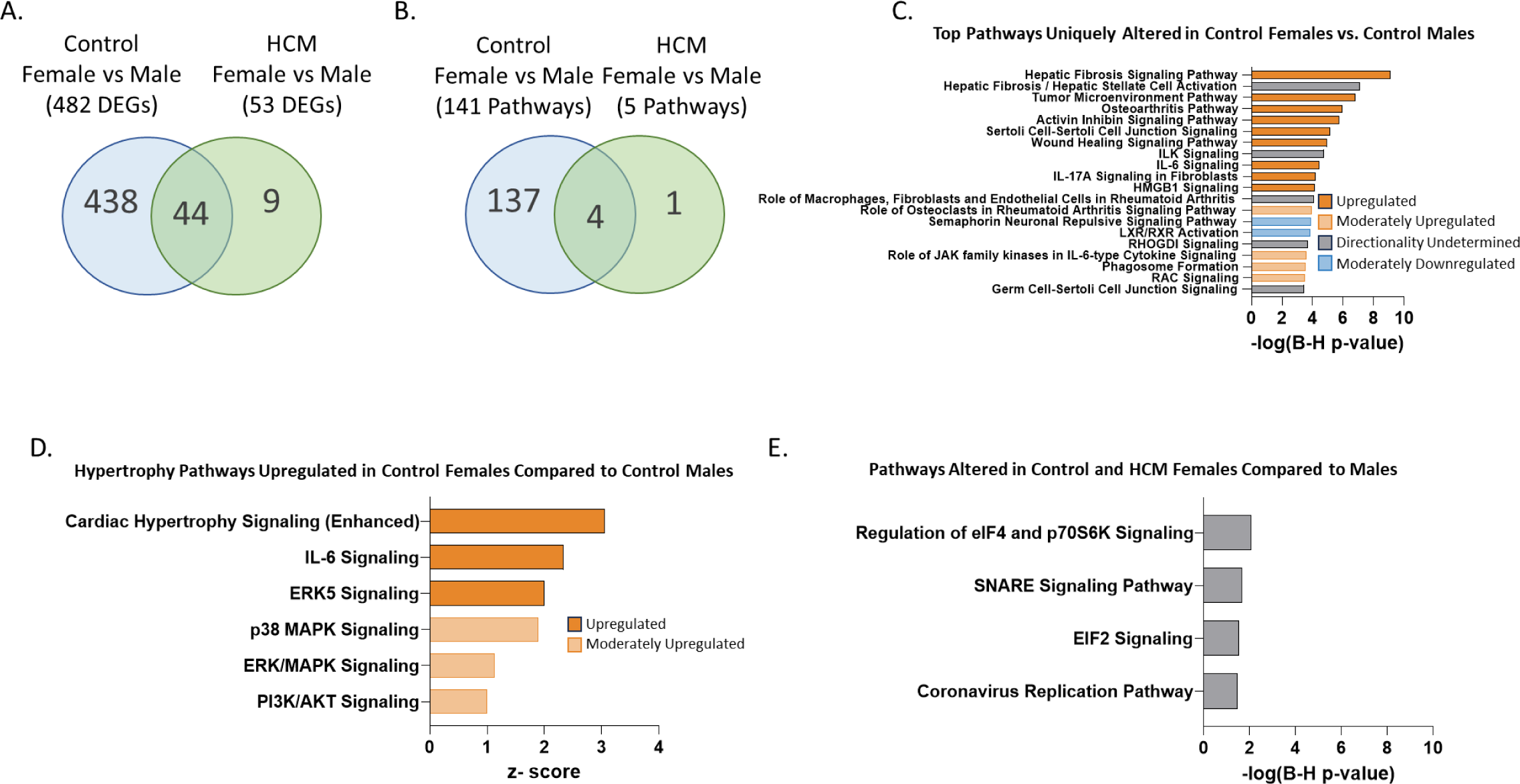
Transcriptome Comparison Between Sexes in Control and HCM Samples. A) Venn diagram of differentially expressed genes (DEGs). B) Venn diagram of differentially regulated pathways. C) Top pathways uniquely altered in control females compared to control males. D) Hypertrophy pathways upregulated in control females compared to control males. E) Pathways altered between sexes in both controls and HCM.

Of the 141 pathways altered between control females and control males, only 4 (3%) were altered between sexes in HCM. There were 137 unique pathways altered in control females compared to control males (**Figure 1B**). The most statistically altered pathways unique to control females vs. control males include fibrotic, immune, and cell adhesion pathways (**Figure 1C**).

Control females had transcriptional upregulation of 6 hypertrophy pathways compared to males, which interestingly were not differentially expressed between sexes in HCM (**Figure 1D**): cardiac hypertrophy signaling (enhanced) (z-score=3.1), IL-6 signaling (z-score=2.3), ERK5 signaling (z-score=2.0), p38 MAPK signaling (z-score=1.9), ERK/MAPK signaling (z-score=1.1), PI3K/AKT signaling (z-score=1.0), and STAT3 signaling (z-score=1). The four sex-specific pathways conserved both when comparing HCM males to HCM females and control males to control females were involved in protein synthesis, cell signaling, and the coronavirus replication pathway (**Figure 1E)**.

### There is More Transcriptional Dysregulation in HCM Females

There were 1983 DEGs between HCM females and control females while there were 1064 DEGs when comparing HCM males to control males; between these 2 comparisons, 802 DEGs were shared (**Figure 2A**). There were 1181 (60%) DEGs uniquely altered in HCM females and 262 (25%) uniquely altered in HCM males. Pathway analysis showed 276 pathways that were altered between HCM females compared to control females (**Figure 2B**). There were 205 pathways altered between HCM males and control males (**Figure 2B**). There were 97 pathways (35%) uniquely altered in HCM females compared to control females, including cytoskeletal signaling pathways, inflammatory, and metabolic pathways (**Figure 2C**). In contrast, there were only 26 uniquely altered pathways (13%) HCM males, including immune pathways, metabolic pathways, and calcium handling (**Figure 2D**). Finally, there were 179 pathways commonly altered in both HCM females and HCM males compared with their respective controls, all the pathways were altered in the same direction, with 121 (68%) being more severely dysregulated in HCM females (**Figure 2E**). Notably, cardiac hypertrophy pathways were transcriptionally downregulated in both HCM females and HCM males, but more strongly downregulated in HCM females for the majority (5 of 6) including cardiac hypertrophy signaling (enhanced) (**Figure 2F**).

**Figure 2.**
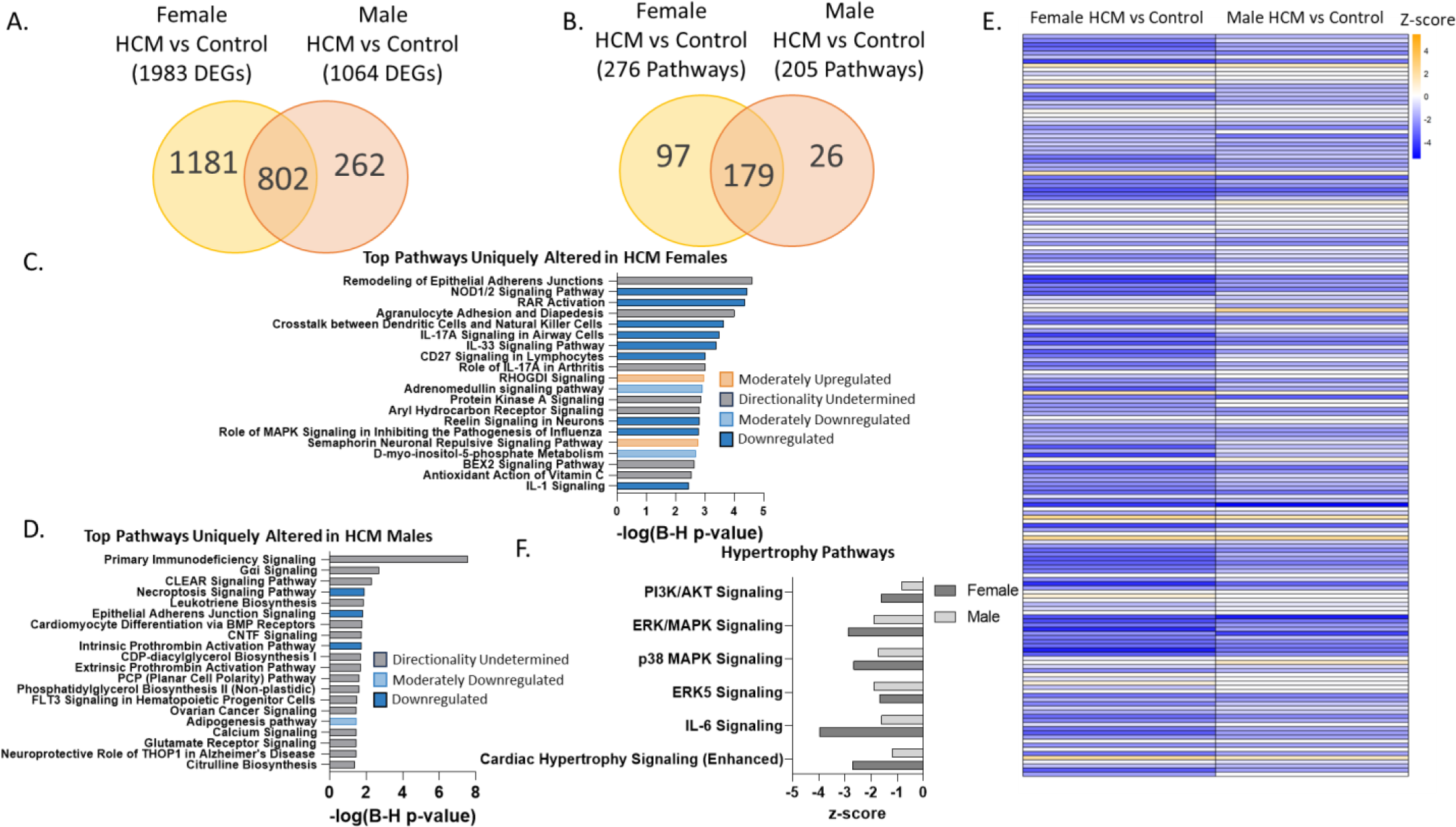
Transcriptome Comparison Between Disease and Control in Females and Males. A) Venn diagram of differentially expressed genes (DEGs). B) Venn diagram of differentially regulated pathways. C) Top pathways uniquely altered in HCM females compared to control females. D) Top pathways uniquely altered in HCM males compared to control males. E) Heatmap of pathways altered in both HCM males and HCM females compared to respective controls. F) Hypertrophy pathways altered in both HCM females and HCM males compared to their respective controls.

### The Proteome of HCM is Similar Between Sexes

In total, 4665 protein were detected in the cardiac samples. PCA plotting showed separation between HCM and controls but not between sexes (**Supplemental Figure 2A**). Similar to the transcriptome, four comparisons were performed: control females vs. control males (**Supplemental Figure 2B**), HCM females vs. HCM males (**Supplemental Figure 2C**), HCM females vs. control females (**Supplemental Figure 2D**), and HCM males compared with control males (**Supplemental Figure 2E**). There were 4 DEPs between control females and control males with 2 being uniquely altered in controls (**Figure 3A**). There were 4 DEPs between HCM females and HCM males with 2 being uniquely altered in HCM (**Figure 3A**). Pathway analysis showed 3 pathways that were commonly altered between sexes in both control and HCM: VEGF, Regulation of eIF4 and p70S6K, and EIF2 signaling (**Figures 3B and 3C**).

**Figure 3.**
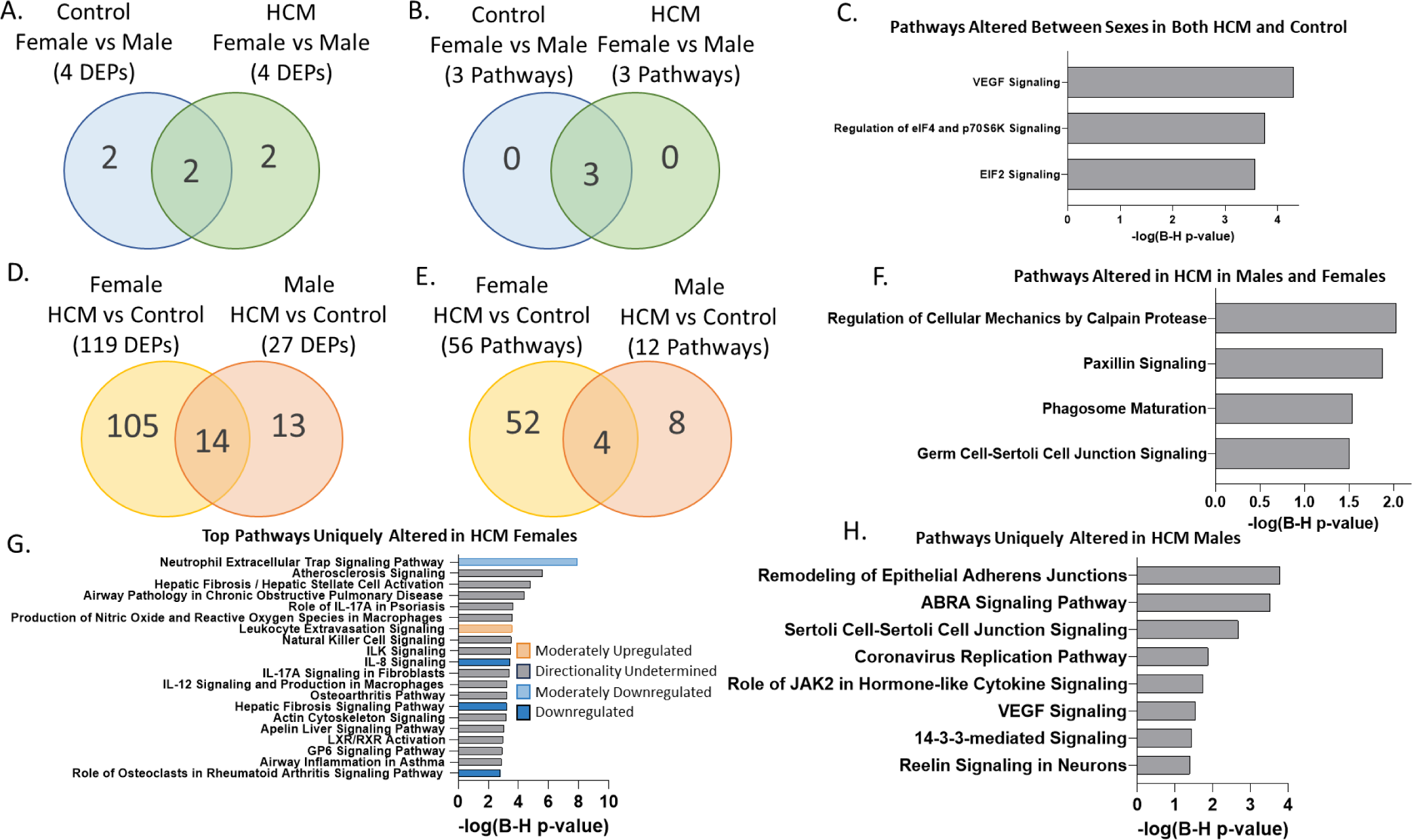
Proteome Comparison. A) Venn diagram of differentially expressed proteins (DEPs) between sexes in control and HCM samples. B) Venn diagram of differentially regulated pathways between sexes in control and HCM samples. C) Pathways altered between sexes in both control and HCM samples. D) Venn diagram of differentially expressed proteins (DEPs) between HCM and control in males and females. E) Venn diagram of differentially regulated pathways between HCM and control in males and females. F) Pathways altered due to HCM in both males and females. G) Top pathways uniquely altered in HCM females. H) Top pathways uniquely altered in HCM males.

### HCM Females Have Increased Dysregulation in the Proteome

There were 119 DEPs between HCM females and control females and 27 DEPs between HCM males and control males (**Figure 3D**). Since the number of samples is the same between both comparisons, any difference in number of DEPs is due to biological differences not due to a difference in power. Only 14 (11%) of the pathways found between HCM females and control females were also altered in HCM males (**Figure 3D**). Pathway analysis showed 56 pathways were altered between HCM females and control females while only 12 pathways were altered between HCM males and control males (**Figure 3E**). Only 4 (7%) of pathways altered in HCM females were also altered in HCM males: regulation of cellular mechanics by calpain protease, paxillin signaling, phagosome maturation, and germ cell-Sertoli junction signaling (**Figure 3F**). There were 52 pathways altered in HCM females but not HCM males including dysregulation of immune/inflammatory pathways including IL-17A signaling, ILK signaling, leukocyte extravasation, and hepatic fibrotic signaling (**Figure 3G**). In addition, two hypertrophy pathway were found to be altered in HCM females but not HCM males: IL-6 and ERK/MAPK signaling. There were 8 pathways uniquely altered in HCM males including remodeling of epithelial adherens junctions, ABRA signaling, VEGF signaling, and 14-3-3-mediated signaling **(Figure 3H**).

### HCM Females Have Increased Dysregulation in the Phosphoproteome

PCA plotting of the phosphoproteome showed clear separation of HCM and controls while there was some separation between male controls and female controls **(Supplemental Figure 3A**).

There was no separation between HCM males and HCM females. Differential phosphorylation analysis was performed comparing control females and control males (**Supplemental Figure 3B**), HCM females and HCM males (**Supplemental Figure 3C**), HCM females and control females (**Supplemental Figure 3D**), and HCM males and control males (**Supplemental Figure 3E**).

Comparing control samples there were 11 DPPs between sexes with 10 (91%) being uniquely altered in controls (**Figure 4A**). Only 1 (9%) DPP was altered between sexes in HCM, showing HCM abolishes sex-specific differences in the phosphoproteome as it does in the transcriptome. There were no unique DPPs when directly comparing HCM males and HCM females (**Figure 4A**). Pathway analysis showed 7 pathways altered between control males and control females; no pathways were altered directly between HCM males and HCM females (**Figure 4B**). The pathways altered in controls included several lipid metabolism and signaling pathways, protein synthesis, and unfolded protein response pathways (**Figure 4C**).

**Figure 4.**
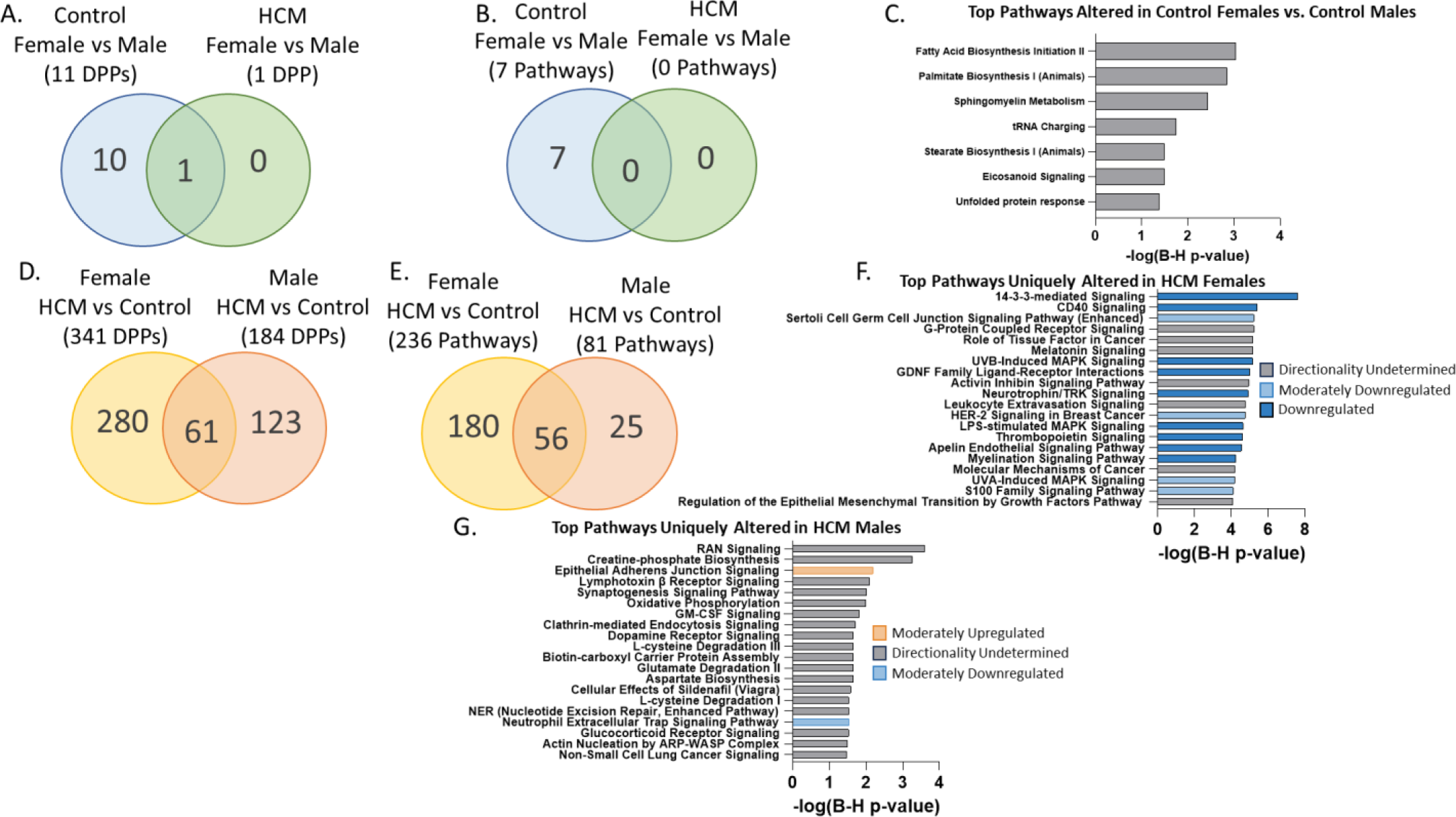
Phosphoproteome Comparison. A) Venn diagram of differentially phosphorylated proteins (DPPs) between sexes in control and HCM samples. B) Venn diagram of differentially phosphorylated pathways between sexes in control and HCM samples. C) Top pathways altered by phosphorylation in control females vs. control males. D) Venn diagram of differentially phosphorylated proteins (DPPs) between HCM and control in males and females. E) Venn diagram of differentially phosphorylated pathways between HCM and control in males and females. F) Top pathways uniquely altered by phosphorylation in HCM females. G) Top pathways uniquely altered in HCM males.

There were 341 DPPs between HCM females and control females, and 184 DPPs between HCM males and control males (**Figure 4D**). Only 61 DPPs were in common between females and males compared to their respective controls: 18% of female DPPs and 33% of male DPPs (**Figure 4D**). Pathway analysis showed dysregulation of 236 pathways in HCM females compared to the control females, and 81 pathways altered in HCM males compared to control males (**Figure 4E**). Only 56 pathways (24%) altered in HCM females compared with control females were also altered in HCM males compared to control males. The top unique pathways altered in HCM females include downregulation of 14-3-3-signaling, CD40 signaling, and several MAPK associated proteins such as UVB-induced MAPK signaling **(Figure 4F).** The top unique pathways altered in HCM males are shown in **Figure 4G**. Interestingly, there was predicted inactivation of 4 established hypertrophy pathways in HCM females compared to control females: ERK/MAPK signaling (z-score = -1.3), IL-6 signaling (z-score = -3.0), p38 MAPK signaling (z-score = -2.4), and cardiac hypertrophy signaling (enhanced) (z-score = -2.8); only one of those pathways was altered in males (ERK/MAPK signaling (z-score = 1.6)), but was found to be activated rather than inactivated (**Figure 5A**). Of the 56 pathways commonly altered in both HCM males and HCM females, 37 (66%) were more strongly altered (larger |z-score|) in females than males (**Figure 5B**). Thus, overall, the phosphoproteome of HCM is more severely dysregulated in females, and we observed inactivation of hypertrophy pathways in HCM females via hypophosphorylation, compared to activation of similar pathways in HCM males via hyperphosphorylation.

**Figure 5.**
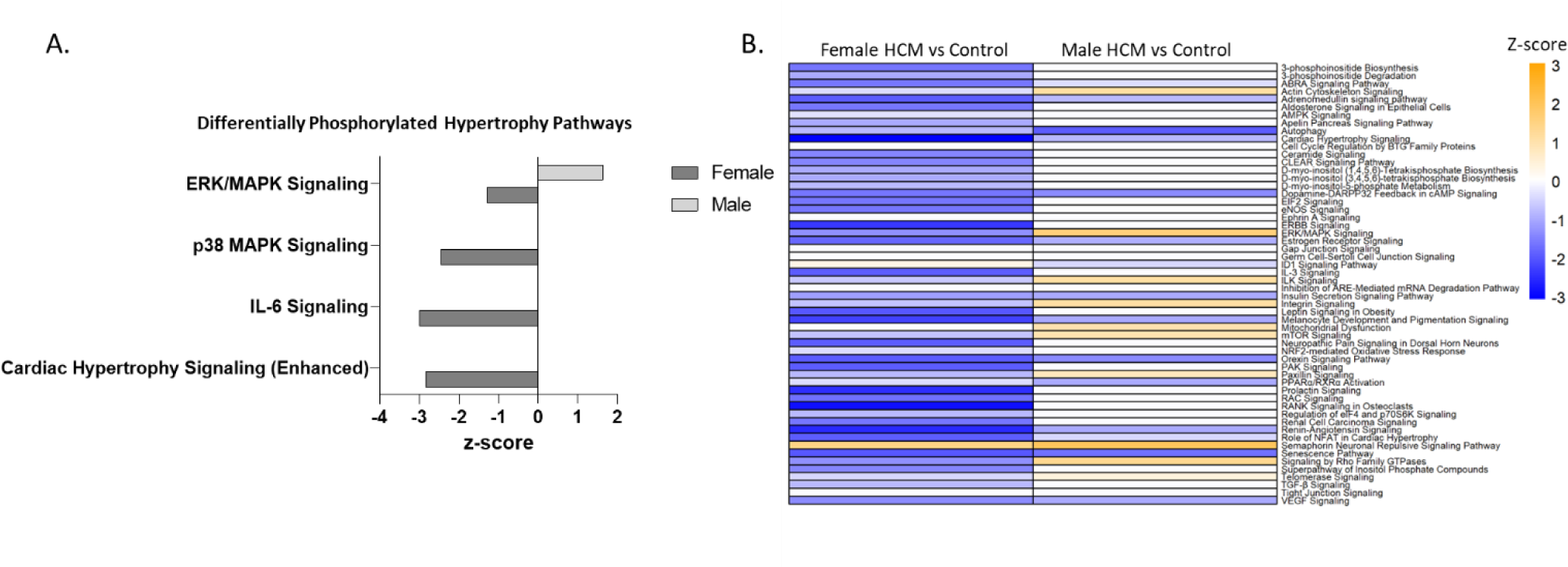
Phosphorylation Comparison Continued. A) Differentially phosphorylated hypertrophy pathways in both HCM females and HCM males compared to their respective controls. B) Heatmap of differentially phosphorylated pathways in HCM females and HCM males compared to their respective controls.

### Summary of Findings

In summary, we found that at baseline, control females have transcriptional upregulation of hypertrophy pathways compared to control males (**Figure 6**). HCM abolishes sex-specific differences found in healthy controls (**Figure 6**). There is transcriptional downregulation of hypertrophy pathways in both HCM males and HCM females, but HCM females have downregulation at the phosphoproteome level while HCM males show activation of these pathways (**Figure 6**). Finally, HCM females have increased dysregulation of the transcriptome and phosphoproteome compared with HCM males (**Figure 6**).

**Figure 6.**
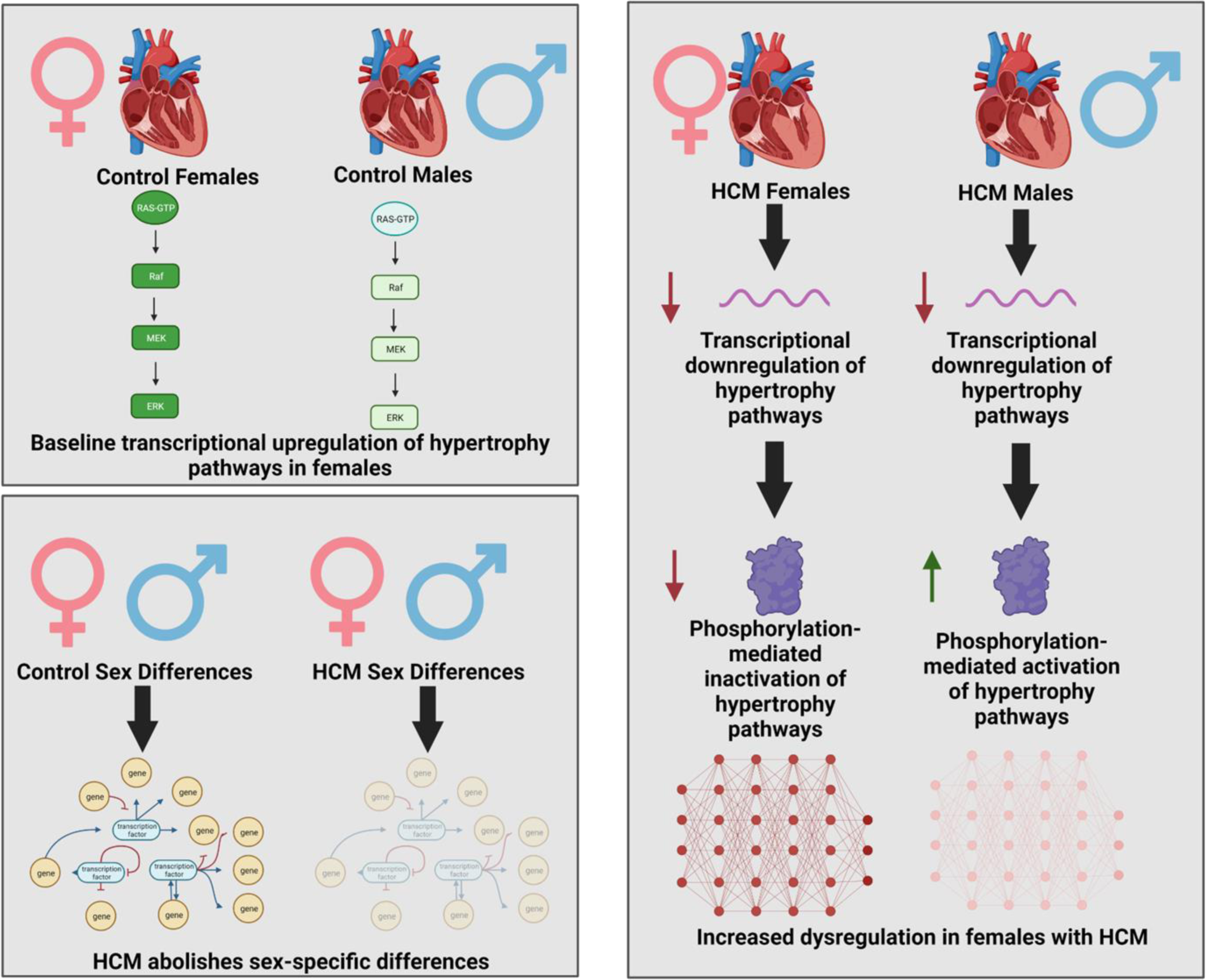
**Summary of Findings.**

## Discussion

### Sex Disparities in Cardiovascular Diseases

Cardiology has notoriously been a field of medicine with clear disparities between males and females, most notably in the misdiagnosis and undertreatment of myocardial infarctions^1, 17^. The disparity is partly due to clinical bias and failure to recognize the unique presentation in women; however, there are also sex-specific biological risk factors for women demonstrating biology plays a critical role^17^. Recent initiatives are in place to try and address the need for better management and understanding of myocardial infarctions in women; however, other areas in cardiology are still in need of improvement.

Specifically, women with HCM are less likely to be diagnosed, but they are more likely to have symptoms, develop heart failure, and have life-threatening arrhythmias^8, 9, 11–14^. There is a paucity of studies investigating the underlying biological mechanisms for these clinical differences. Cellular studies of cardiomyocytes from patients with HCM have shown increased fibrosis and increased compliant titin between males and females^18^. In addition, another study performed proteomic analysis of myectomy tissue from sarcomere-positive HCM patients and found 46 differentially expressed proteins with increased levels of tubulin and heat shock proteins in females^19^. To our knowledge, this manuscript is the first multi-omics comparison of sex differences in HCM patients, including both sarcomere-positive and genotype-negative samples, to identify a universal sex-specific HCM disease signature and provide an atlas of transcriptional, proteomic, and phosphoproteomic differences in HCM.

### HCM Females Have Increased Dysregulation

Overall, we found a general, common profile in the transcriptome, proteome, and phosphoproteome between males and females with HCM as evidenced by a lack of separation on PCA plotting. A direct comparison between HCM males and HCM females did elucidate small, direct differences in the transcriptome and (phospho)proteome. Indirectly, at every level of the transcriptome and (phospho)proteome, HCM females showed a higher number of alterations compared to female controls suggesting increased transcriptional and proteomic dysregulation.

In addition, for the majority of pathways in both the transcriptome and phosphoproteome, the degree of dysregulation was greater in females compared to males.

Additionally, we observed a transcriptional downregulation of pathways such as ERK5, p38 MAPK, ERK/MAPK, IL-6, and cardiac hypertrophy, which have previously been implicated in cardiac hypertrophy^20–23^, in HCM females and HCM males compared to their respective controls. However, the degree of downregulation was much greater for females than males for 5 of 6 pathways. Recently, we demonstrated HCM myectomy samples were characterized by transcriptional downregulation of hypertrophy pathways, possibly representing a counter-regulatory attempt to curb further hypertrophy^15^. Thus, in this stage of HCM it appears females had even greater counter-regulatory downregulation of hypertrophy pathways.

### HCM Abolishes Normal Sex-Specific Transcriptional Differences

While previous studies have shown there are clear transcriptional differences in the myocardium of females and males^24, 25^, we found almost 500 DEGs between control females and control males of which interestingly only 10% were found to be different between HCM males and females highlighting HCM transcriptional differences are abolished between sexes. Consistent with previous studies, we found differences in transcriptional expression of many inflammatory pathways between control males and females^26^ with females showing transcriptional upregulation of many innate and adaptive immunity pathways. Differences in expression of inflammatory and immune pathways have been proposed as partly responsible for biological differences in cardiovascular outcomes between males and females^26, 27^. Whether inflammatory pathways play a role in sex-specific differences in HCM requires further studies.

One novel and interesting observation was the transcriptional upregulation of several hypertrophy pathways in control females compared to male controls, including cardiac hypertrophy (enhanced), IL-6, ERK5, p38 MAPK, ERK/MAPK, and PI3K/AKT signaling. This increased basal level of pathway activation compared to males could play a part in the differences in disease pathogenesis and disease susceptibility. Further studies are necessary to determine the functional consequences of this observation and the temporal transcriptional and phosphoproteomic differences between sexes during the course of disease. Although there were fewer changes in the phosphoproteome due to smaller sample sizes for the phosphoproteomics study, 11 DPPs were identified between control females and males, while only 1 (9%) DPP was found between HCM females and HCM males.

### Differences in Phosphorylation are Found Between Females and Males

Previously, we identified in HCM myectomy tissue that there is hyperphosphorylation and activation of ERK5 and ERK/MAPK signaling^15^. Interestingly, here we found females with HCM had hypophosphorylation and inactivation of several cardiac hypertrophy pathways including cardiac hypertrophy signaling (enhanced), p38 MAPK signaling, and IL-6 signaling, while. in contrast in males, ERK/MAPK signaling was hyperphosphorylated and activated similar to previously published studies suggesting RAS/MAPK signaling plays a role in the cardiac hypertrophy in obstructive HCM^15, 28^.

### Temporal Model of Hypertrophy Pathways in HCM

Taken together, this multi-omics atlas can be used to develop and further investigate sex-specific models for hypertrophy pathways in HCM (**Figure 7**). Studies previously demonstrated activation of hypertrophy pathways in other forms of cardiac hypertrophy is initiated by transcriptional activation^29^. Previously, for obstructive HCM, we demonstrated **translational** activation of hypertrophy pathways, but **transcriptional** downregulation of hypertrophy pathways which we speculated to be a counter-regulatory attempt to curb further hypertrophy^15^.

**Figure 7.**
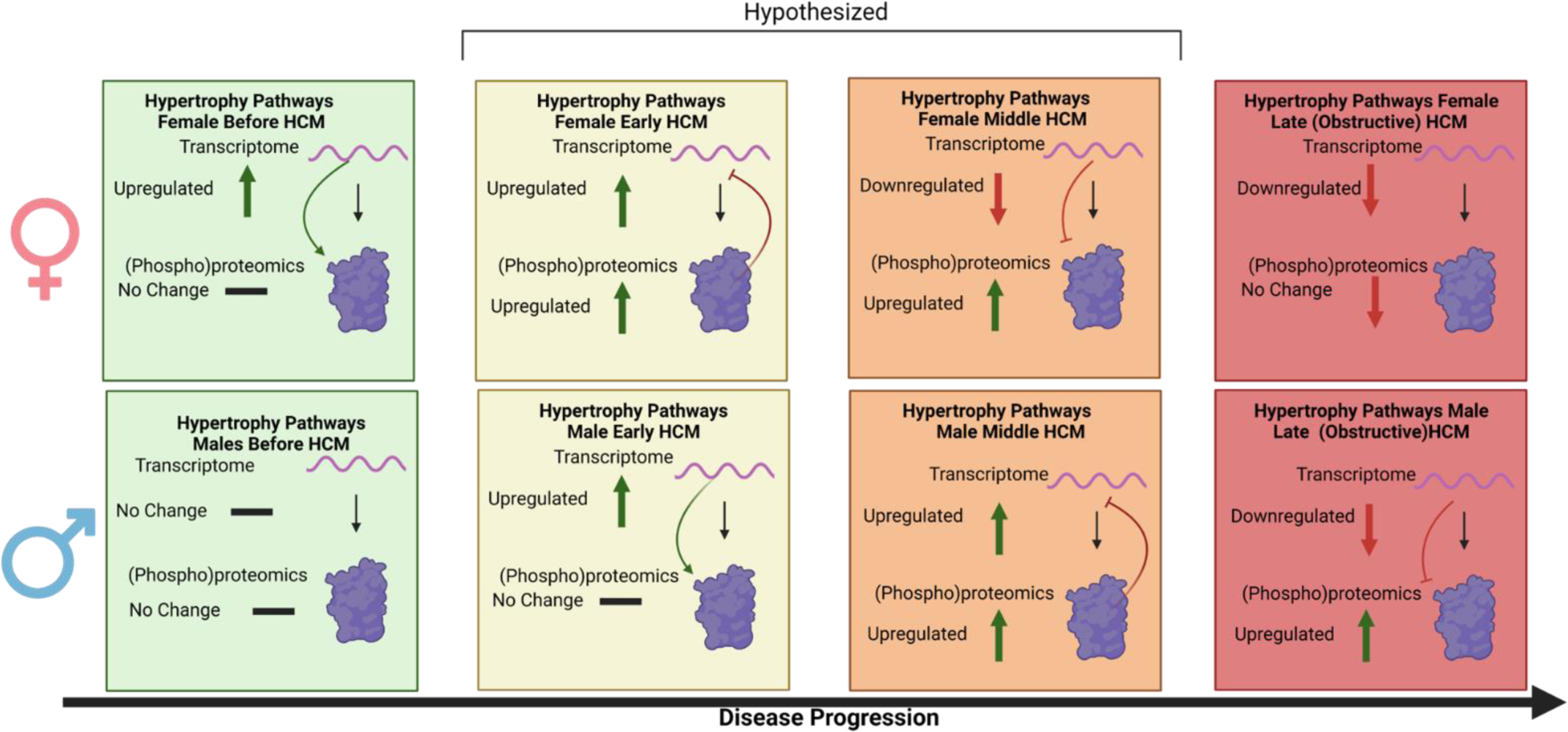
**Proposed Sex-Specific Temporal Model of Hypertrophy Pathways in Hypertrophic Cardiomyopathy (HCM).**

Thus, we herein posit HCM pathobiology starts with transcriptional activation of hypertrophy pathways leading to activation at the protein level, but with crucial sex-specific, temporal differences. In fact, our data shows that HCM females have both transcriptional and post-translational downregulation of hypertrophy pathways, while males still have active pathways at phosphoproteome level. As such, it appears that females progress more rapidly towards having downregulation of hypertrophy through this negative feedback loop, possibly as a result of a higher activity of hypertrophy-associated pathways before disease (i.e. in controls). Temporal studies are necessary to compare males and females with HCM and observe when the differences arise and functionally characterize the impact of these changes.

## Conclusion

Myectomy samples from patients with obstructive HCM have a similar mult-omics profile between males and females; however, there are subtle differences between males and females which may underly differences in clinical outcomes between sexes. Additionally, HCM appears to abolish sex-specific differences found in controls.

## Acknowledgements

We would like to thank the Mayo Clinic Genome Analysis Core and the Mayo Clinic Proteomics Core for their assistance in acquiring rigorous and high-quality data.

