## Supplemental Figures for "A Multi-Omics Atlas of Sex-Specific Differences in Obstructive Hypertrophic Cardiomyopathy"

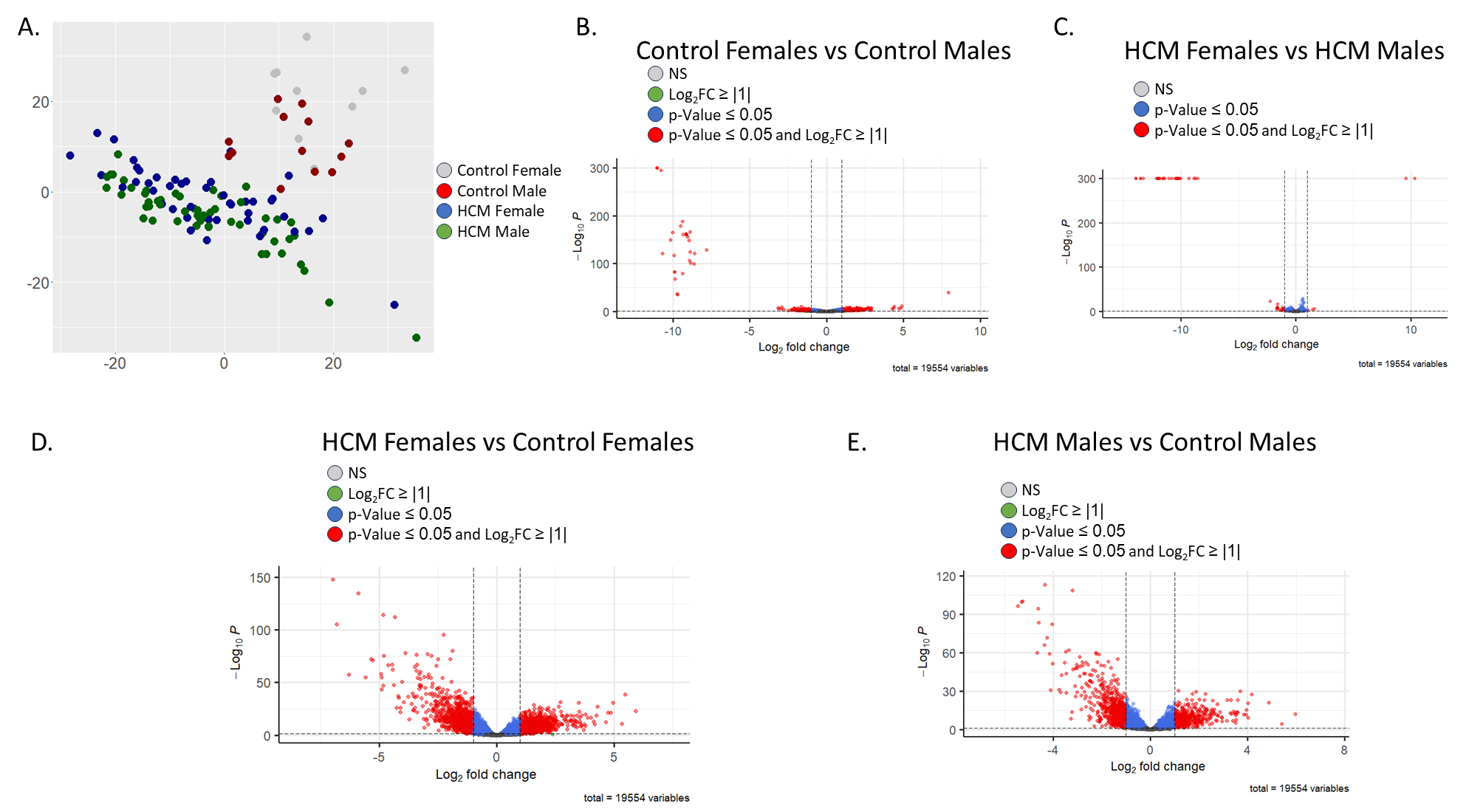

**Supplemental Figure 1. Transcriptional Analysis**. A) Log normalized PCA plot of transcriptome. B) Volcano plot of differential expression analysis control females vs. males. C) Volcano plot of differential expression analysis HCM females vs. males. C) Volcano plot of differential expression analysis females with HCM vs. female controls. D) Volcano plot of differential expression analysis males with HCM vs. male controls. FC, fold change; NS, not significant.

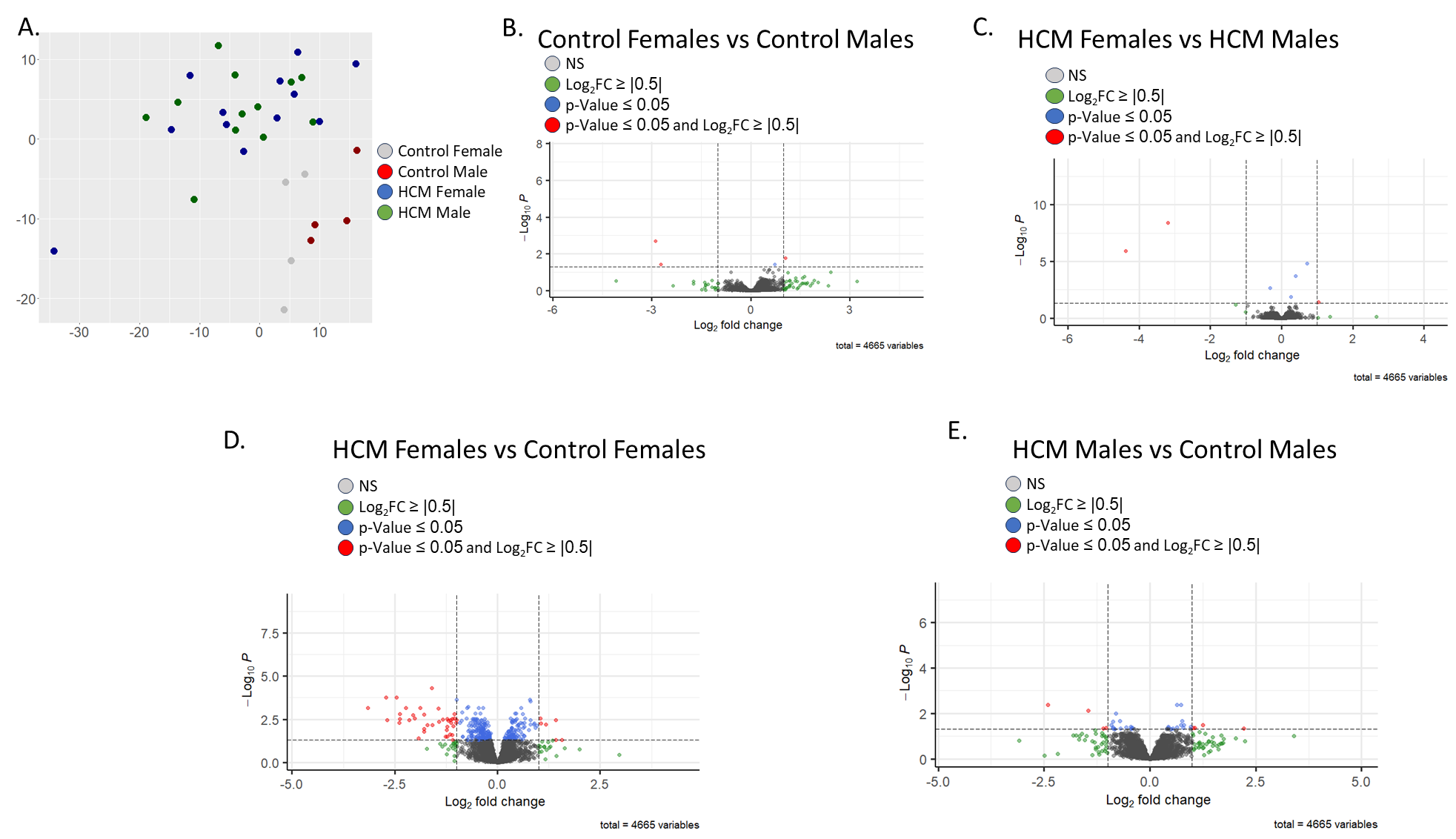

**Supplemental Figure 2. Proteome Analysis**. A) PCA plot of proteome. B) Volcano plot of differential expression analysis control females vs. males. C) Volcano plot of differential expression analysis HCM females vs. males. C) Volcano plot of differential expression analysis females with HCM vs. female controls. D) Volcano plot of differential expression analysis males with HCM vs. male controls. FC, fold change; NS, not significant.

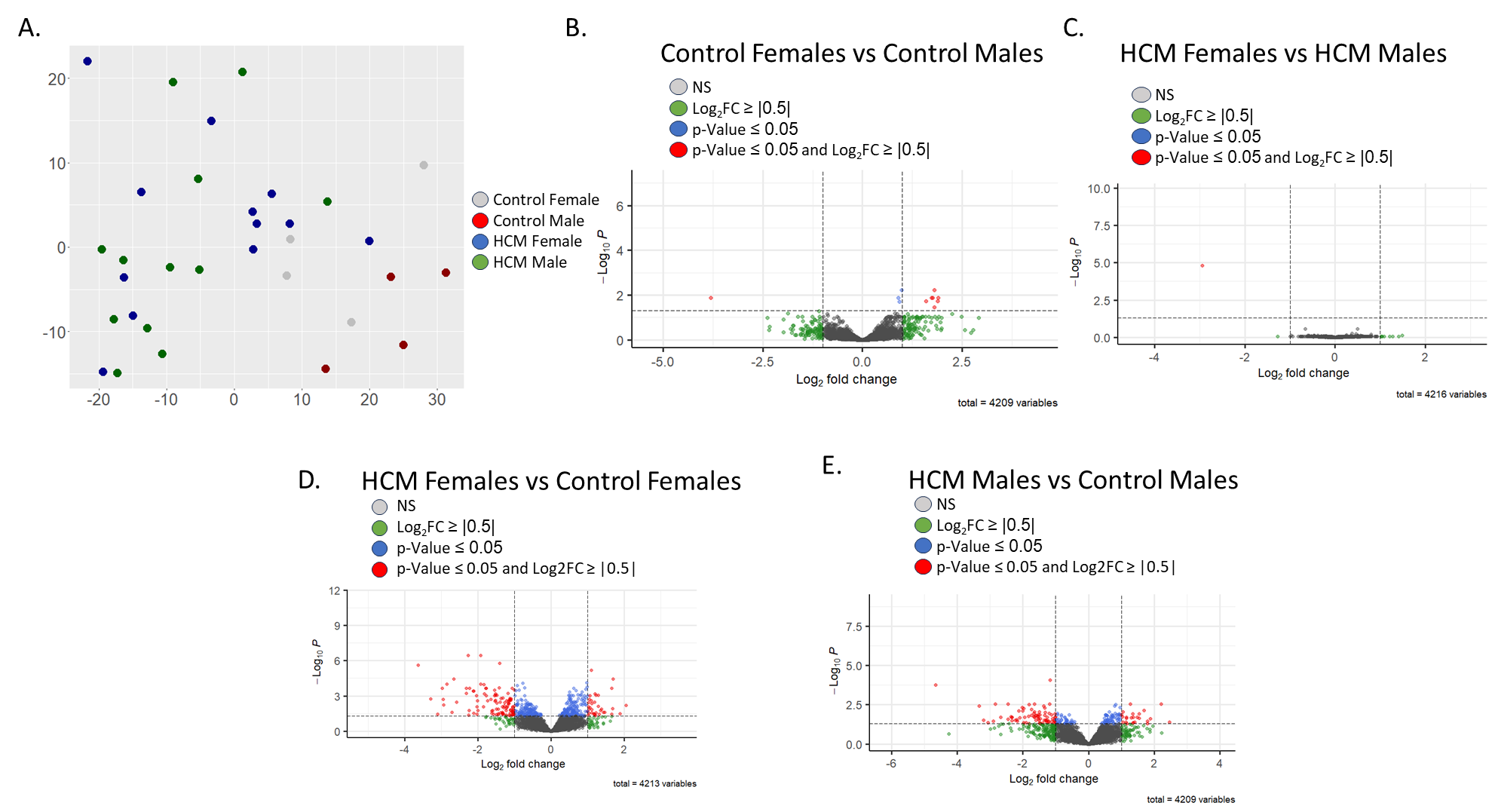

**Supplemental Figure 3. Phosphoproteome Analysis**. A) PCA plot of phosphoproteome. B) Volcano plot of differential phosphorylation analysis control females vs. males. C) Volcano plot of differential phosphorylation analysis HCM females vs. males. C) Volcano plot of differential phosphorylation analysis females with HCM vs. female controls. D) Volcano plot of differential phosphorylation analysis males with HCM vs. male controls. FC, fold change; NS, not significant.
