## Supplemental Table 1 for "A Multi-Omics Atlas of Sex-Specific Differences in Obstructive Hypertrophic Cardiomyopathy"

|  | HCM Females (n=44) | HCM Males (n=53) | p-value |
| --- | --- | --- | --- |
| Average age at diagnosis, y | 43 ± 20 | 39 ± 19 | 0.3 |
| Average age of myectomy, y | 48 ± 19 | 46 ± 18 | 0.5 |
| Family Hx of HCM, n (%) | 15 (15) | 19 (20) | 0.9 |
| Family Hx SCA, n (%) | 8 (8) | 6 (6) | 0.3 |
| SCA, n (%) | 0 (0) | 1 (1) | 0.9 |
| ICD, n (%) | 10 (10) | 17 (18) | 0.3 |
| Max LVWT at diastole, mm | 22 ± 6 | 24 ± 7 | 0.3 |
| LV mass index, g/m^2 | 168 ± 65 | 205 ± 85 | 0.03 |
| Max LVOT gradient, mm Hg | 89 ± 42 | 84 ± 38 | 0.5 |

**Supplemental Table 1. Demographic Comparison Males and Females.** HCM, hypertrophic cardiomyopathy; HX, history; ICD, implantable cardioverter defibrillator; LV, left ventricle; LVOT, left ventricular outflow tract; LVWT, left ventricular wall thickness; Max, maximum; SCA, sudden cardiac arrest.
